## Supplemental Information for "Single domain antibodies against enteric pathogen virulence factors are active as curli fiber fusions on probiotic *E. coli* Nissle 1917"

### Supporting Information

#### Table S1: Strains and plasmids

| (a) Strains | | |
|---|---|---|
| **Strain** | **Description/comments** | **Reference** |
| Mach1 | *E.coli*, str.K-12 $F^-$ ϕ80(*lac*Z) ΔM15 Δ*lac*X74 *hsd*R($rK^- mK^+$) Δ*recA*1398 *end*A1 *ton*A, cloning strain | Thermo Fisher Scientific |
| MC1061 | *E. coli*, str. K-12 $F^-$ $\lambda^-$ Δ(ara-leu)7697 [araD139]B/r Δ(codB-lacI)3 galK16 galE15 $e14^-$ mcrA0 relA1 rpsL150($Str^R$) spoT1 mcrB1 hsdR2($r^-m^+$) | Casadaban *et al.*, 1980 [93] |
| PBP8 | *E. coli* Nissle 1917, Δ*csgBACDEFG*::*$Cm^R$* | Praveschotinunt *et al.*, 2018 [52] |
| 2457T | *Shigella flexneri* 2a strain 2457T | Mills *et al.*, 1992 [94] |
| E2348/69 | *E. coli* strain O127:H6/**EPEC** ; widely used as a model for EPEC infection | Iguchi *et al.*, 2009 [95] |
| EDL933 | *E. coli* strain O157:H7/**EHEC**, *stx1 stx2 eae espP/pssA* $hly_{EHEC}$) | O'Brien *et al.*, 1983 [96] |
| E22 | *E. coli* strain O103:K-H2/**REPEC**, *rha-* | Camguilhem *et al.*, 1989 [97] |
| 3014-2 | *E. coli* strain O153:H-/**REHEC**, sorbitol+ | García *et al.*, 2002 [98] |
| DBS770 | *Citrobacter rodentium*, derivative of strain ICC168, chloramphenicol resistant. Lysogenized with Stx2dact-producing phage ϕ1720. Produces Shiga toxin. | Mallick *et al.*, 2012 [99] |
| DBS771 | *Citrobacter rodentium*, chloramphenical resistant, kanamycin resistant, *stx2dact*. DBS770 with KanR cassette inserted into prophage *stx* genes. Does not produce Shiga toxin. | Mallick *et al.*, 2012 [99] |
| E10 | *E. coli* strain O119:H6/EPEC wt | Girón *et al*., 2002 [51] |

| **(b) Plasmids** | | |
|---|---|---|
| **Plasmid** | **Description/comments** | **Reference** |
| pL6FO | CsgA-VHH expression vector, contains synthetic curli operon *csgBACEFG* under an IPTG-inducible promoter. | Kan *et al.*, 2019 [88] |
| pGex-2t | Vector for EHEC intimin GST-Int465 (*eaeA*, EDL933 (C-terminal 465aa, bp1636-3082) (Yu and Kaper, 1992 [100])); EPEC GST-TirM (E2348/69 TirM 120aa) (Burland *et al.*, 1998 [101]); REPEC GST-TirM (E22 TirM 120aa) | Pharmacia, Liu *et al.*, 1999 [50] |
| pDEST15 | Vector for *C. rodentium* intimin GST-Int400C (*eaeA*, *Citrobacter rodentium* strain DBS100 (C-terminal 400aa)); | Thermo Fisher Scientific |
| pGex-4t2 | Vector for EPEC intimin (*eaeA*, E2348/69 (C-terminal 400aa)); REPEC intimin (*eaeA*, E22 (C-terminal 400aa)) | Pharmacia |
| pET15b | Vector for EPEC His-TirM (E2348/69 TirM 120aa) (Burland *et al.*, 1998 [101]) | Novagen |
| pDest17 | Vector for REPEC His-TirM (E22 TirM 120aa); *C. rodentium* His-TirM (DBS100 TirM 120aa) | Thermo Fisher Scientific |
| pMalc2 | Vector for EHEC intimin MBP-Int395 (*eaeA*, EDL933 (C-terminal 395aa, bp1618-2082) (Liu *et al.*, 2002 [102])) | New England Biolabs, Liu *et al.*, 1999 [50] |
| pUC19 | Vector for intimin expression in MC1061 (*eaeA*, full-length, EDL933 bp -30-3011; *eaeA*, full-length, JPN15 bp 1614-4537) (Yu and Kaper, 1992 [100]) | Yanisch-Perron *et al.*, 1985 [103] |

**Table S2: Aligned sequences of VHHs binding pathogenic *E. coli* virulence factors**. Complementarity determining regions (CDR1, 2 and 3) are highlighted and appear in order from left to right.

Anti-Fla:

JUV-B11: SGGGLAQPGGSLRLSCTSTGHT-LDDYAIGWFRQAPGKERERVACASASGI-TTNYADSVKGRFTISRDKAKNMVYLQMNSLQPEDTAVYYCAA---TPYYGDVCVRAAFESRGQGTQLTVSS

JUV-C4: TGGGLVQAGGSLTLSCVASGRA-VSSFAMGWFRQIPGREQRDFVAFIGDYGLTTYYANSVKGRFTISRSNAENTLYLQMNSLEFEDAAVYFCAA--RDAYRSGTTNPSAYDYWGQGTQVTVSS

JUV-E8: SGGGLVQAGGSLRLSCAASGRT-SSTYTMGWFRQAPGKEREFAAAIRSSGS-GTYYADSVKGRFTISRDGAKNTVYLQMNSLKPEDTAVYYCAA--RGNPIYSVYDVRTYDLWGQGTQVTVSS

JUV-G8: TGGGLVQPGGSLRLSCAASGSI-VSFNAMVWYREAPGKQREWVAQITPSSK--TMYKDSVKGRFTISSDNAKNMVYLQMNNLKPEDTAVYYCNGD-------------RGVAWGPGTQVIVSS

JUV-H5: SGGGLVQPGGSLRLSCAASEMS-FSIKRMGWFREAPGKPREWVAQITPAGS--TNYAETVKGRFTISRDNAKNTVYLQMNSLKPDDTAVYYCNT-------------LPGIAWGQGTQVIVSL

JWU-F3: SGGGLVQPGGSLRLTCVSSLSD-FRLTNMAWYRQTPGSERDVVAGISPNGI--TSYHASVQDRFNISRDNARKTLFLQMNSLKPEDSGVYYCNI-----------RWGSLLPWGQGTQVTVSP

JWU-H4: TGGGSVQPGGSLRLSCAPSGFS-LAYYGVGWFRQAPGKEREALACISRFGD-DTYYADSAKGRFTVSRDNAKNTVYLEMNNLKPEDTGVYYCAAGWVVVTDESCGSGDAYNYWGRGTQVTVSS

JXE-B1: TGGGLVQAGDSLRLSCAASSGRNFSNYATGWFRQAPGKEQEFVASISRSGR-STYYADSAKGRFTISRDNARNTVYLQMNSLKPEDTADYYCAAHETQWPNGLGWVRGFYDLWGQGTQVTVSS

Anti-intimin:

JWS-H4: SGGGLVQAGGSLRLSCTTSASIFSGYRMGWFRQAPGKQREFVASIADG-QNTPYADSVMGRFTISRDNAKNTVYLQMSNLKPEDTAIYYCKS-----------------WGYTDWGQGTQVTVSS

JWT-C1: S-GGLVQPGGSLRLSCAASGFTLANSAIGWFRQAPGKGREAVSCISTSAGTTNYASSVKGRFTTARDNAKNMAYLQMDNLKSGDTGVYECAA------GWSIDCSGYILPAADVWGQGTQVTVSS

JWU-D8: TGGGLVQPGGSLRLSCAASGFYFSGYWMHWVRQVQGQGLKWVSGINIDDTKSSYTDSVKGRFTISRDNTKNTLYLQMDSLQPEDTGVYYCAR-------DRRAGQISGGYDPDYRGQGTQVTVSS

JWU-G8: SGGGLVQPGGSLRLSCATSGFYFAGYWMHWVRQVPGQGLEWVSGIDLGSTMLNYRDSVKGRFSISRDNAKNTVYLQMHSLKPEDTALYFCAR-------DRRAGATSGGYDPDVRGQGTQVTVSS

JXN-E2: T-GGLVPPGGSLQLSCTSSGFFLDYLGVAWFRQAPGNEREGVSCIDYTGDNIAYASSMKGRATISRDKDANTVTLEMNGLKPEDTAVYYCAAHRSATTYADGKYRCPLENEYDYWGQGIQVTVSS

Anti-Tir:

JVB-C6: TGGGLVQPGGSLTLSCAASGFSITENAMGWARQVPGKGLEWVSLV---------YSGGNTYYAESIEGRFTISRDNAKNTVYLRMTSLKPEDTGVYYCAA----REAVRVAGPPADVWGQGTQVTVSS

JVB-G4: SGGGLVEAGGSLRLSCSASGRASGDGHLGWFGDGALAWFRQAPGKEREYVAAVGHSGTDTYYSDSVKGRFTISRDNAKSMGYLQMDNLRPDDTGIYYCAL------DLHLGQPGYDYWGQGTQVIVSS

JVB-G8: TGGGLVQAGGSLRLSCVASGFTFNDYVLTWFRQAPGKGREGVASI--------SPAFGNTFYADSVKGRFTITSDSAKKQVCLQMNNLKSEDTAVYFCAADPTVNFGPVLRAENYRHWGQGTQVTVSS

JVC-C6: SGGGLVQAGGSLRLSCAASGVTFVDYATGWFRQAPGKEREGIACI--------SNGASGSFVVNSVKGRFTTSTDNAKNTAYLQMNDLKPEDTATYYCAL-LGRTGGHCTDPNEYSYWGQGTQVTVSS

JVC-D10: T-GGLVQTGDSLTLSCVVSGRGFCDWAMAWFRQAPGKEREKVSAI--------GWSGQDTYYSEPAKGRFTISRDNAKNTVWLRMTNLKSEDTAVYYCAA-------ATRAYADYDYWGQGTQVTVSS

JVC-E5: TGGGTVQAGSSLRLSCTASGFTFTDAAMGWFRQAPGKEREFVAAI--------NWDSANKYYADSVKGRFTISRDNAKNTVYLEMTALKPEDTADYICAG--DAKLGHVATSDVWRFWGQGTQVTVSS

JVA-A1: S-GGSVEAGDSLRLSCAASGRGFGDGALAWFRQAPGKEREYVAAV--------GHSGTDTYYADSVKGRFTISRDNAKNMGYLQMDSLRPDDTGVYYCAL------DLHLGQPGYDYWGQGTQVTVSS

JVA-C8: SGGGLVQAGGSLRLSCAASGFTFVDYGIAWVRQAPGKEREGVSCV--------STSNRSQWYADSVKGRFTISSDNAKNTVYLQMDNLKPEDTAVYYCTR----IDVGDNCRDGGGYRGQGTQVTISS

JVA-C9: TGGGLVQAGGSLRLSCVDSGRTFGDNAMGWFRQAPGKEREYVAAI--------GYNGDSTYYLNSVKGRFTISRDNAKNTLYLQMNSLQLEDTGVYICAA------KTRVTTKEYDFWGQGTQVTVSS

JVA-D4: TGGGLAQAGGSLRLSCATSGFTFADYAIGWFRQAPGKEREGVACI--------STSDNGQYSADSVKGRFTISKDNAKKTVYLQMNSLKPDDTAIYYCHL--IDVGSNCRKGKDDGFWGQGTQVTVSS

JVA-F6: SGGGLVQAGDSLRLSCVTSGRSFSEAAMGWFRQAPGKERELMTSI--------GWNGDRTYYADSVKGRFTISRDNAKNTVYLQMNGLTPNDTAVYYCAA-------AVRASKGYEYWGQGTQVTVSS

JVA-D11: SGGGLVQAGGSLRLSCAASGFTFNDYAKAWFRRAPGKEREGISAI--------SSMGESTFYADSVKGRFTISSDSAKNTVYLQMNSLKPEDTAVYYCAADPTVKWGHVLREENYDYWGQGTQVTVVS

JVA-E10: SGGGLVQAGGSLRLSCVASGRTFSDYAMGWFRQAPGKEREFVAAI--------SWNGGSTYYADSVKGRFTISRDNAKNTMYLQMNSLKSEDTAVYYCIS-EGRGLRQVKTATDWEYWGQGTQVTVSP

JVA-G1: SGGGLAQAGGSLKLSCVASGFTFADYAIGWFRQAPGKEREGVACI--------SNSVGSTYYSDSVKGRVTISSDNVKKTVYLQMNSLKPEDTAFYYCAL--IDHGMNCRNSNGIHYWGKGTLVTVSS

Anti-EspA:

JXF-D7: SGGGLVQTGGSLRLSCSASGFALEYYAVGWFRQAPGKEREGVSCF--SGSDGSKFHAQFVKGRFTISLDKEKNTVDLTMNNLKPEDTAVYYCAVA-GPSDHQCDLGMTWYHRWGQGTQVTVSS

JYB-B1: TGGGLVQAGGSLRLSCAISGFSLGDYAIGWFRQAPEKEREGVAFS--GSLGNTY-YPDSMKGRFTISRDDAENAVYLEMKNLKPEDTAVYRCTS----GGTFDAVVLGLSTYWGQGTQVTVSS

JYB-B8: TGGGLVQPGGSLRLSCAASGFTFSTYWMYWVRQAPGKGLEWVSTI--DTGGSDTYYADSVKGRFTISRDNVKNTLYLQMDNLKPEDTALYYCGS---SRDIVIVTTLRDFDYWGQGTQVTVFS

JYB-D1: TGGGLVQAGGSLRLSCAASGNTFSYTAMAWFRQAPGKQRELVARI--SSGRGPTKYADSVKGRFTISRDNTLNTVWLQMNDLKPEDTAVYFCNT----LKYSGESSYIAGDSWGQGTQVTVSS

JYB-H4: SGGGLVQPGGSLRLSCAASGVTLEYYAIGWFRQVPGKEREGVSCI--STSGAGTNYADSVKGRFTISKDNAKNTVYLQMNSLKPEDTAVYYCAA-RDFTEVPIGGCEWEYDYWGQGTQVTVSS

JYB-H6: TGGGLVQPGGSLRLSCVASGFILDAYTIGWFRQAPGKEREGVASI--NGSGFSTNYADSVKGRFTISRDNAKNTVWLQMNSLKPEDTAVYYCAA--ALGLLTPRLESTPFDDWGQGTQVTVSS

JYF-D8: SGGALVQPGGSLRLSCAASGINLDYYAIGWFRQAPGKERKGVSCI-SHVDDDRIYYSDSVKGRFTISRDNAKNTVYLQMNSLEPEDTAVYYCATA-GPSDYPCDLELWTYRRWGQGTQVTVSS

JXF-H9: SGGGLAQTGGSLRLSCAASGFRLEYYAVGWFRQAPGKEREGVSCL--SGSDGSTYNAEFAKGRFTISRDDAKNTVYLLMNSLKPEDTAVYYCAVA-GPSDYQCDLGTRWYHRWGQGTVVTVSA

JXF-C4: SGGGLVQAGGSLRLSCVAYGRTANTFKEYAMGWFRHNPGEDHEFVGGISQNGDEAYFDDSVKGRFTPSRDNAKNTMYLQMNSLRPEDTAAYYCVAQRSSERLVGDMYSAMDSWGKGTLVTVST

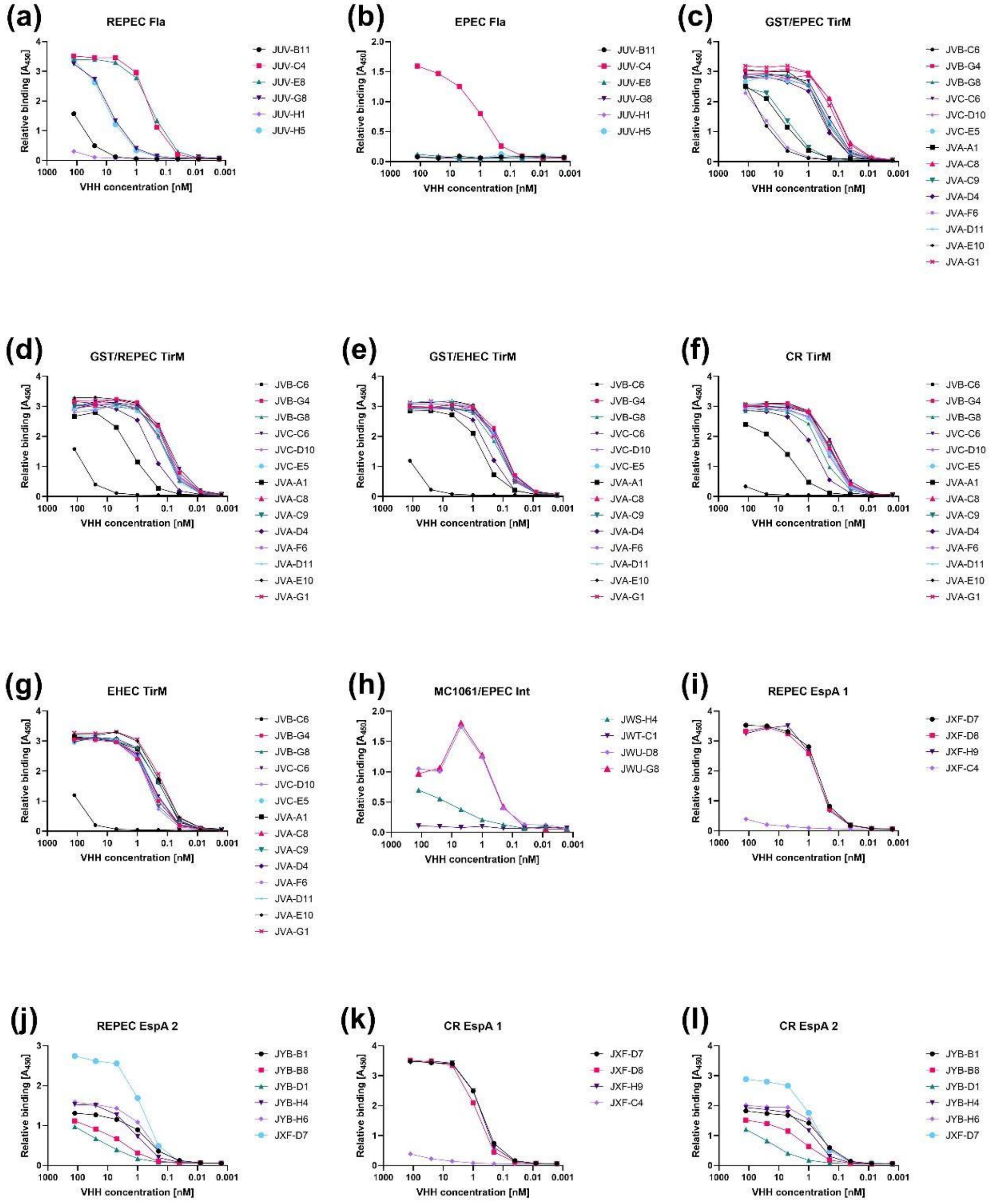


**Figure S1: Assessing the binding properties of selected VHHs via ELISA.** Antigens used were homologues of Fla (a-b), Tir (c-g), Int (h) or EspA (i-l) corresponding to either EPEC (b, c, h), EHEC (e, g), REPEC (a, d, i, j) or *C. rodentium* (f, k, l). In each assay, antigen was either directly added to the plate in purified form (a-b, e-g, i-l), bound to the plate by an adsorbed noncompeting VHH (c-d), or displayed on the surface of MC1061 (h).

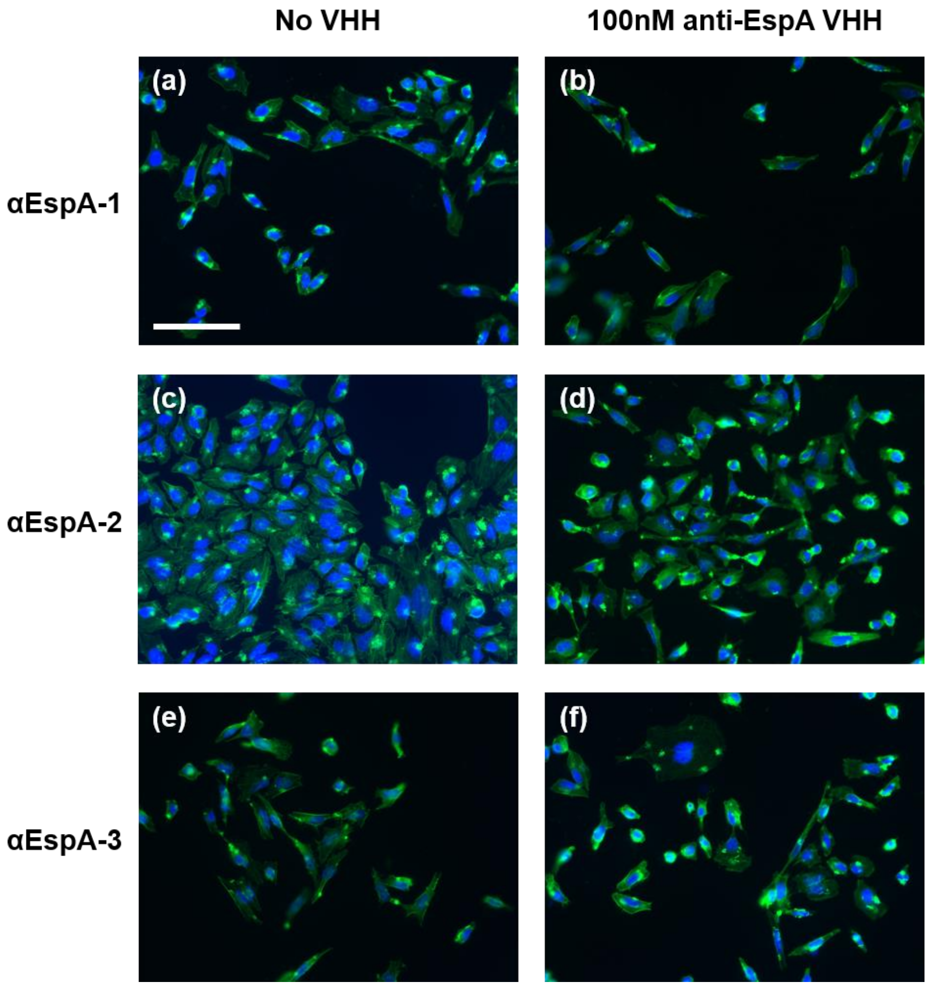


**Figure S2: Anti-EspA VHHs did not inhibit pedestal formation.** HeLa cells were exposed to EPEC incubated with VHH, fixed and stained with DAPI (blue) and Alexa Fluor-488 Phalloidin (green). Similar to the "no VHH" negative control (a, c, e), all anti-EspA VHHs tested (b, d, f) resulted in the formation of pedestals (though not all anti EspA-VHHs were tested) (scale bar = 100 μm).

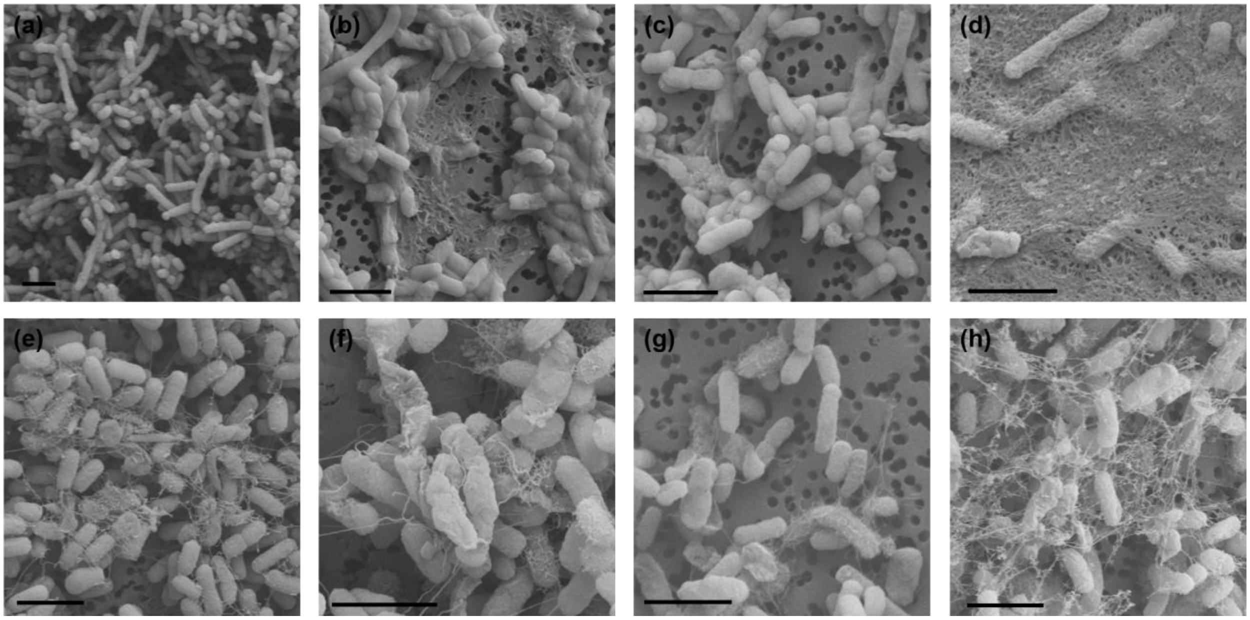


**Figure S3: Representative FESEM images of EcN expressing CsgA and CsgA-VHH.** (a) EcN (PBP8) with no plasmid, expressing no curli fibers. (b-h) EcN expressing CsgA-VHH, exhibiting a range of fiber morphologies. (b) CsgA-αStx2, (c) CsgA-αInt-12, (d) CsgA-αInt-17, (e) CsgA-αFla-3, (f) CsgA-αFla-4, (g) CsgA-αIpaD-1, (h) CsgA-αgp900-2 (scale bar = 2 μm).

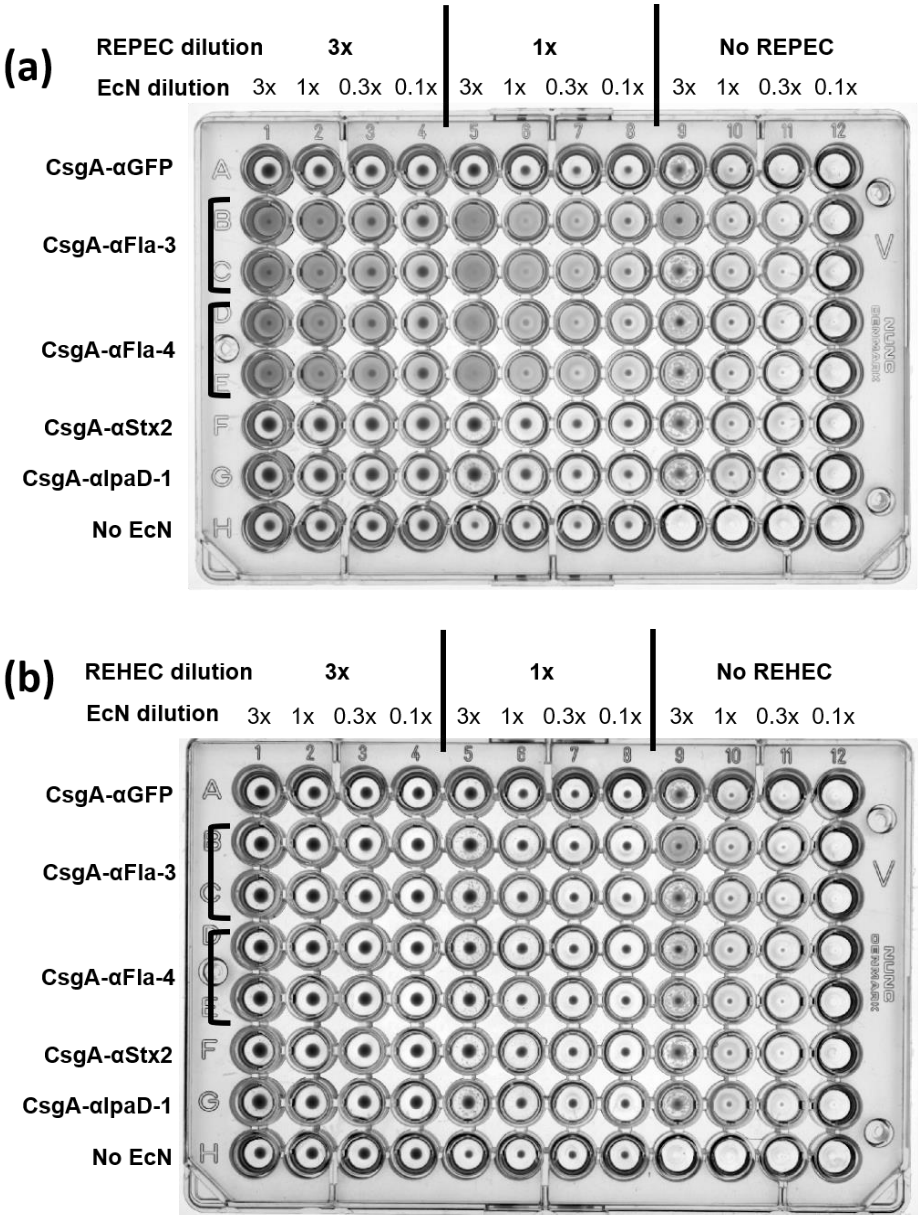


**Figure S4: EcN expressing CsgA-αFla can induce REPEC aggregation.** Suspensions of EcN expressing CsgA-VHH were mixed with either REPEC (a) or REHEC (b) and allowed to settle overnight in conical 96-well plates. Aggregation was only observed when REPEC was mixed with CsgA-αFla-3 and -4.

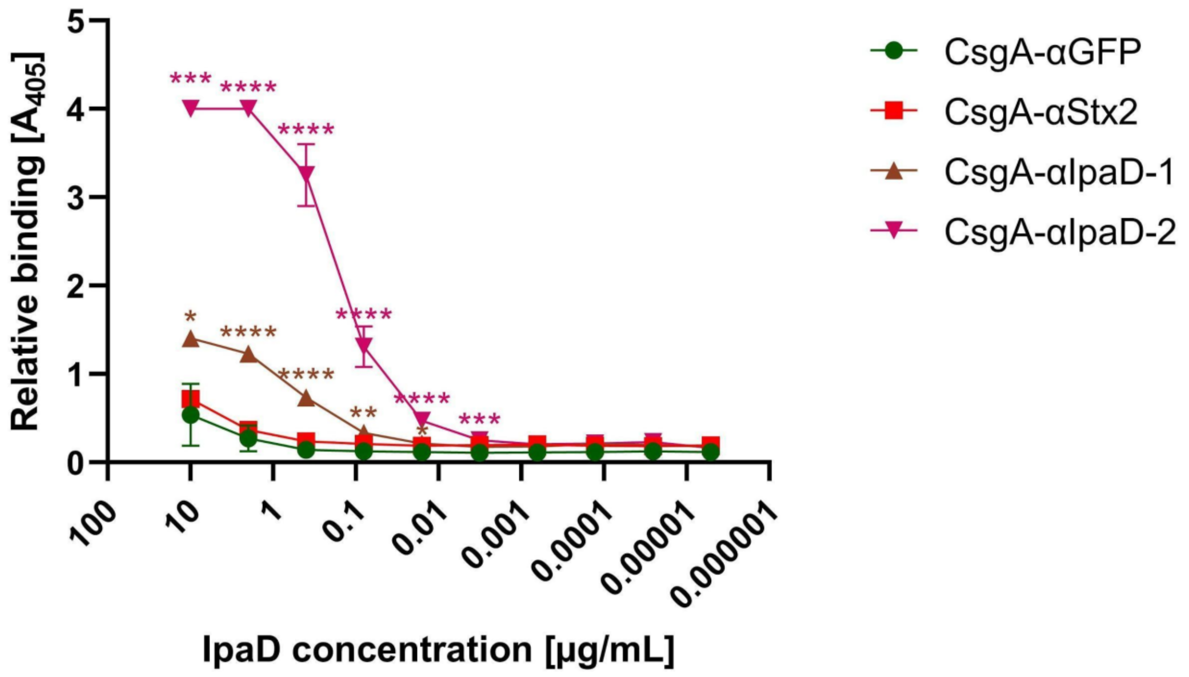


**Figure S5: EcN expressing CsgA-αIpaD can bind soluble IpaD.** ELISA demonstrated the ability of CsgA-αIpaD to bind soluble IpaD. EcN was adsorbed onto a well plate, followed by incubation with varying IpaD concentrations. Binding of IpaD to the adsorbed EcN was then detected by a specific non-competing VHH (JMK-H2, Barta *et, al.*, 2017 [20]), followed by an anti-Etag IgG-HRP conjugate. EcN expressing either CsgA-αIpaD-1 or CsgA-αIpaD-2 significantly outperformed the off-target negative control (CsgA-αGFP). Data presented as mean ± SD. Two-way ANOVA ($P < 0.0001$) was performed to test the presence of difference between conditions, P-values calculated by Welch's t-test. * $P < 0.05$; ** $P < 0.01$; *** $P < 0.001$; **** $P < 0.0001$.

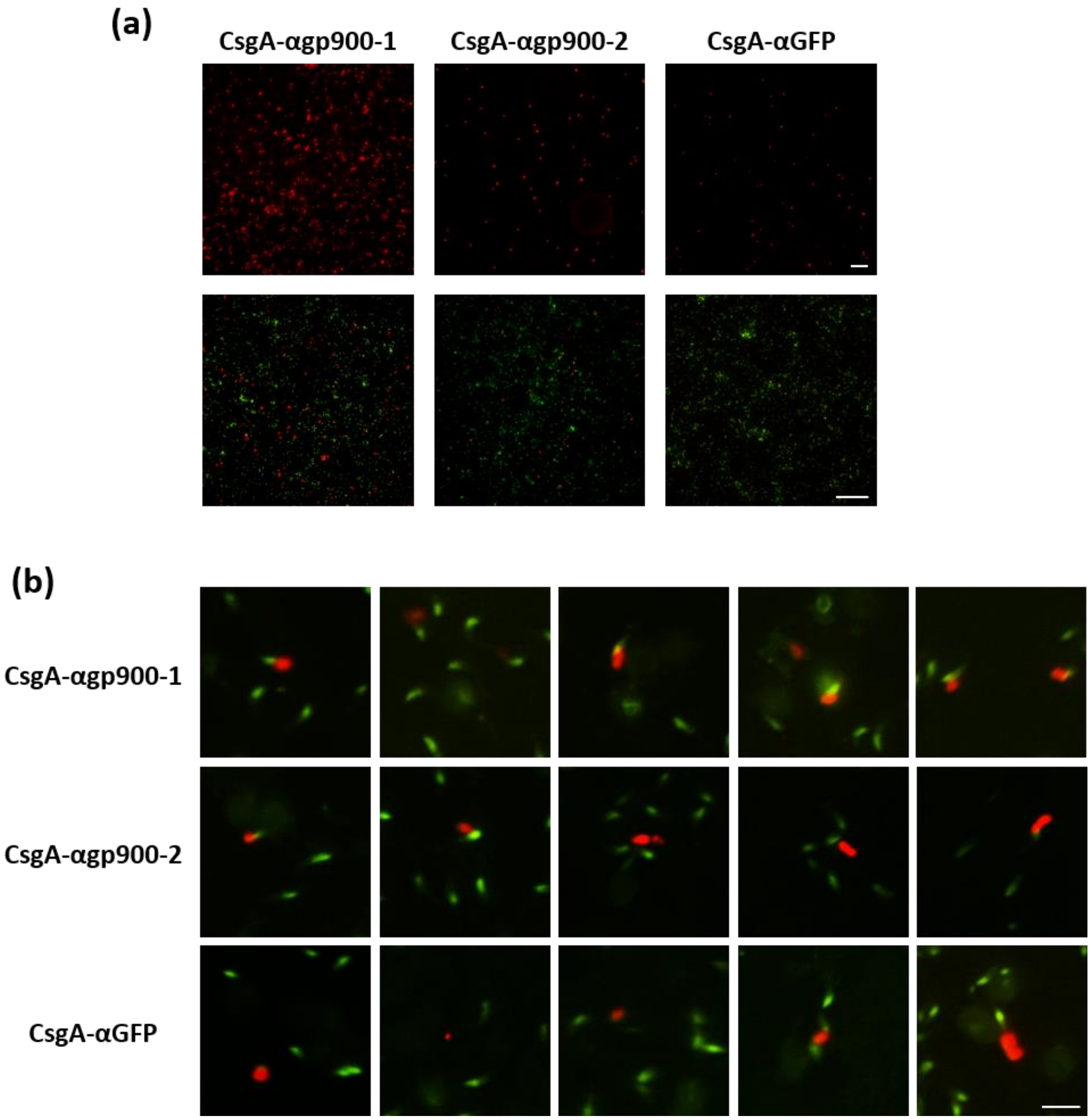


**Figure S6: EcN expressing CsgA-αgp900 exhibit increased attachment to and colocalization with *C. parvum* sporozoites.** (a) Fluorescent micrographs demonstrating increased attachment of EcN (red) to *C. parvum* sporozoites, counterstained in green in the bottom panels (scale bar = 50 μm). (b) While EcN (red) expressing CsgA-αgp900-1 (and to a lesser extent CsgA-αgp900-2) consistently colocalize with sporozoites (green), the CsgA-αGFP negative control was often observed away from the green fluorescent foci, consistent with nonspecific binding (scale bar = 5 μm).
